# The onset and offset of noxious stimuli robustly modulate perceived pain intensity

**DOI:** 10.1101/2020.03.18.996769

**Authors:** Benedict J. Alter, Mya Sandi Aung, Irina A. Strigo, Howard L. Fields

## Abstract

Reported pain intensity depends not only on stimulus intensity but also on previously experienced pain. A painfully hot temperature applied to the skin evokes a lower subjective pain intensity if immediately preceded by a higher temperature, a phenomenon called offset analgesia. This is typically evoked using a three-step noxious heat stimulus. In other clinical and laboratory settings, prior pain experience may increase pain intensity as well. Therefore, we hypothesized that even small increases in stimulus intensity within the noxious range would be accompanied by enhanced reported pain intensity. To test this possibility, we inverted the intensity order of the three-step stimulus, so that the same hot temperature is immediately preceded by an increase from a transiently lowered temperature. Using healthy volunteer subjects, we observed a disproportionate increase in pain intensity during the novel, inverted, three-step stimulus. This disproportionate increase is similar in magnitude to that of offset analgesia. Control stimuli demonstrate that these changes in pain intensity are distinct from habituation. The magnitudes of offset analgesia and the disproportionate increase in pain intensity correlate with each other but not with the absolute noxious stimulus temperature. These observations suggest that the disproportionate increase in pain intensity represents an “onset hyperalgesia.” Finally, the magnitude of both offset analgesia and onset hyperalgesia depends on preceding temperature changes. Overall, this study finds that perceptual enhancement of noxious stimulus change occurs bidirectionally and that this depends on the intensity and direction of change of the immediately preceding stimulus.

## Introduction

Controlled psychophysical studies using acute stimuli reveal a consistent relationship between stimulus intensity and reported subjective pain intensity [1]. This relationship reflects the activity of the peripheral and central nociceptive transmission pathways that mediate pain perception. However, both Alter, et al. Onset hyperalgesia - pg. 2 clinical and experimental studies have demonstrated that there can be a dramatic dissociation between subjective pain reports and the intensity of the applied nociceptive input. Reported pain is consistently and predictably modifiable by prior experience and expectation, such that equivalent noxious input can yield divergent pain reports when sensory cues associated with lower or higher pain are presented [2, 3]. Placebo analgesia and the increase in pain with nocebo are examples of this process [4, 5]. Importantly, there is a dynamic interaction between the intensity of ongoing pain perception and the impact of superimposed noxious stimulation [6]. Although much progress has been made in understanding how learned predictive cues influence subsequent pain perception, less is known about how changes in nociceptive input can act as predictive cues and dynamically modify the perceptual impact of subsequent noxious input.

In an effort to better understand dynamically changing pain perception, and specifically pain relief, Coghill and others made a significant advance when they discovered and provided detailed studies of what they labeled “offset analgesia” [7, 8]. Using a cutaneous thermode they found that when there is an ongoing stable moderately painful heat stimulus, a one-degree Celsius increase in thermode temperature for 5 seconds followed by a return to the prior noxious temperature elicits a disproportionate drop in pain intensity rating compared with a constant stimulus at the initial temperature – so-called “offset analgesia”. This effect was also observed when the transient increase and decrease were applied on the contralateral arm consistent with a primarily central nervous system mechanism for offset analgesia [9]. Functional imaging studies indicate that offset analgesia is mediated by brainstem pain modulatory circuitry [10–13]. Curiously, although these modulatory circuits are known to exert bidirectional control over nociceptive transmission, no disproportionately large increase in pain with the onset of the higher temperature was reported. Subsequently, elegant modeling of pain intensity with supra-threshold heat stimuli predicted perceptual enhancement of temperature increases as well as decreases[14], although this was not empirically demonstrated. Increases in radiant heat using infra-red light applied to the skin does elicit increases in pain intensity, but again these were less in magnitude than temperature decreases[15].

To clarify the apparent contradiction between reports of offset analgesia without equivalent enhancement of temperature increases and the known bidirectional effects of descending modulation, we re-examined the effect of small changes of stimulus intensity on subsequent pain reports. In the current study, we report that robust onset hyperalgesia exists, consistent with bidirectional perceptual modulation dependent upon the direction of change of the immediately preceding noxious stimulus.

## Materials and Methods

### Subjects

35 female and 39 male subjects signed written informed consent for the study, which was approved by the University of California, San Francisco (UCSF) Institutional Review Board. Subjects were recruited by public notice, including on-line advertising through Craigslist and paper advertising at UCSF. Inclusion criteria were healthy subjects aged 18-50 years old. Exclusion criteria included current significant medical comorbidities requiring frequent medical follow-up, pregnancy, chronic pain, ongoing acute pain, depression, anxiety, bipolar disorder, psychotic disorder, concurrent analgesic use, concurrent psychoactive medications (e.g. benzodiazepines, antidepressants), blindness, deafness, non-fluency in English. For consistency with ongoing studies, exclusion criteria also included inability to undergo magnetic resonance imaging (MRI) for any reason (e.g. non-compatible implants, claustrophobia).

### Testing protocol

The study timeline included recruitment and administration of standardized surveys followed by a single study visit lasting 2-3 hours. During the study visit, subjects completed surveys and then underwent heat pain threshold testing, heat stimulus calibration, and supra-threshold heat pain testing.

The study visit occurred in a research lab located in a clinical building on a hospital campus. A single office room was used. Experimenters included authors B.A. and S.A. All subjects were monitored using a 3-lead electrocardiogram (ECG) which was in place during testing. Breaks were allowed if the subject requested them. Subjects were compensated monetarily for their time and travel.

### Questionnaires

Upon study inclusion, subjects completed electronic (REDCap) or paper surveys. For most subjects, surveys were done electronically prior to study visit. If the electronic survey was not done or incomplete on the day of the study visit, a paper survey was done. The survey packet included self-reported basic demographic information, medical histories, and measures of social status (BSMSS), depression (BDI-II), anxiety (STAI Y-1 and Y-2), impulsivity (BIS-11) and pain catastrophizing (PCS). The BSMSS generates a single ordinal score reflecting the respondent’s education and occupation as well as the education and occupation of their parents and spouse[16]. The BDI-II measures cognitive-affective and somatic-vegetative aspects of depression with excellent psychometric properties across different populations[17]. The STAI measures both state (Y-1) and trait (Y-2) anxiety, producing an ordinal score reflecting apprehension, tension, nervousness, and arousal[18, 19]. The PCS measures catastrophic thinking associated with pain incorporating magnification of pain-related symptoms, rumination about pain, feelings of helplessness, and pessimism about pain-related outcomes [20]. The BIS-11 measures attentional, motor, and non-planning impulsiveness [21] which are associated with reward processing relevant to pain and addiction [22, 23]. After sensory testing outlined below, the STAI Y-2 and the situational pain catastrophizing scale (SPCS), measuring catastrophizing related to a pain experience [24], were administered.

### Equipment and pain reporting

Subjects were seated in a comfortable office chair in front of a desk. All heat stimuli were applied with the Pathway NeuroSensory Analyzer (Medoc; Ramat Yishai, Israel) using an fMRI-compatible, 3×3 cm ATS thermode. Subjects reported heat pain threshold with a button press using the Pathway Patient Response Unit. Subjects reported pain intensity in real time using a Computerized Visual Analogue Scale (COVAS, Medoc) consisting of a 100-mm visual analogue scale anchored by “no pain sensation” on the left and “most intense pain sensation imaginable” on the right. Subjects positioned a slider on this scale to reflect their pain intensity rating in that moment. Slider position over time was recorded using Medoc software.

### Cutaneous heat stimulation

All heat stimuli were applied to the volar surface of the non-dominant forearm. Three locations were used for heat stimulation, rotating from proximal to distal back to proximal. The time between heat stimuli was at least 120 seconds. With site rotation, the time between stimulating the same area of skin was at least 6 minutes. The thermode was placed on the volar forearm to allow full contact with the skin without excessive pressure and secured with a Velcro strap. The thermode was held at 32°C between heat stimuli.

### Heat pain threshold testing

Heat pain threshold was first determined using the method of limits. From a baseline of 32°C, the thermode was warmed at a rate of 1.5 C°/sec until subjects reported the transition from heat to pain via button press. The temperature reached was recorded as the heat pain threshold. The language used to describe the transition was “whenever the sensation changed from heat to pain.” Maximum cutoff temperature was set to 55 °C. Heat pain thresholds from the three skin locations were averaged. This temperature was then used for heat stimulus calibration.

### Heat stimulus calibration

This portion of the protocol established an individually calibrated temperature to elicit a pain rating of ~50 mm on the COVAS at the end of a 30-second constant heat stimulus. The stimulus started at a baseline of 32°C, increased at a rate of 1.5 C°/sec, maintained a constant target temperature for 30 seconds, and returned to baseline temperature at a rate of 6 C°/sec. Subjects were asked to rate their pain on the COVAS in real time during each stimulus. The initial target temperature was chosen based on heat pain threshold. If the threshold was greater than or equal to 45°C, the initial target temperature was the heat threshold temperature. With lower thresholds, the initial target temperature was 2 C° higher than threshold. The stimulus target temperature was increased by 1 C° until pain report was between 40-60 mm on the COVAS. If pain report was greater than 90 mm, the subsequent stimulus was adjusted down by 1 C°. Finally, the temperature producing a pain report of ~50 mm on the COVAS was used for subsequent procedures and is referred to as T1. One C° higher was defined as T2.

### Supra-threshold heat pain testing and pain intensity curve analysis

Subjects rated pain on the COVAS in response to a series of complex and control supra-threshold stimuli (Fig 1A). These included the novel Inverted (Inv) stimulus which started at a baseline of 32°C, maintained at T2 for 5 seconds (t1 period), decreased by 1 C° to T1 for 5 seconds (t2 period), increased by 1 C° (back to T2) for 20 seconds (t3 period), and returned to baseline (Fig 1A&B). Initial rise rate from baseline was 1.5 C°/sec. Rates of change after achieving T2 were 6 C°/sec. The three-step stimulus typically used to measure offset analgesia was included (3step; rise rate 1.5 C°/sec, T1 for 5 seconds, T2 for 5 seconds, T1 for 20 seconds; rates of change and fall rate 6 C°/sec; Fig 1A&C) as well as a novel two-step stimulus (2step; rise rate 1.5 C°/sec, 10 seconds at T2, 20 seconds at T1, rates of change and fall rate 6 C°/sec; Fig 1A&D). Constant control stimuli included a simple step to T1 (T1; 30 seconds at T1, rise rate 1.5 C°/sec, fall rate 6 C°/sec; Fig 1A,C,D) and a simple step to T2 (T2; 30 seconds at T2, rise rate 1.5 C°/sec, fall rate 6 C°/sec; Fig 1A&B).

**Fig 1:**
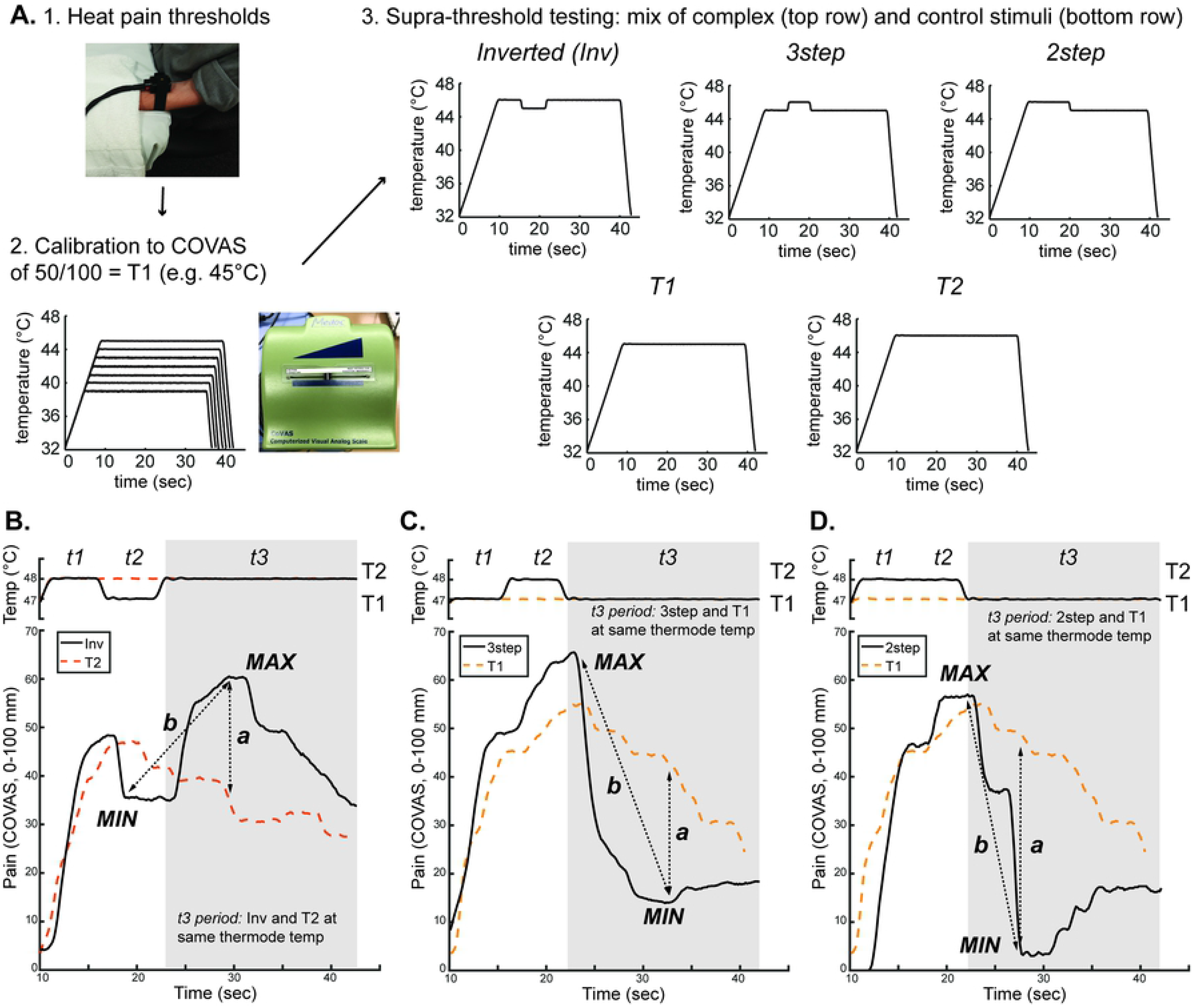
Experimental design and examples of data extraction. **A.** Subjects underwent (1) heat pain threshold testing, (2) an ascending series of suprathreshold, constant, 30-second, temperature stimuli to determine an individualized temperature that would elicit a COVAS pain rating of 50 mm/100 mm, and finally (3) a randomized mixture of suprathreshold, 30-second temperature stimuli. The mixture included novel complex supra-threshold stimuli (top row) and control constant stimuli (bottom row) shown. The temperature curves plotted are examples in which T1 = 45°C. **B.-D.** Examples of continuous pain intensity measured by COVAS and thermode temperature during complex stimuli with appropriate control stimuli superimposed from a single subject. Inverted (Inv) data are plotted with the control T2 stimulus data to measure differences in pain intensity during the *t3* period despite the same thermode temperature (**B.**). 3step (**C.**) and 2step (**D.**) data are plotted with T1 data to measure differences in pain intensity during the *t3* period despite the same thermode temperature. The dotted arrows labeled “*a*” depict measured differences between complex and control curves. For example in **B.**, the *a* arrow shows the difference between the local maximum of the COVAS pain curve during the Inv stimulus and the pain intensity at the same timepoint during the control T2 stimulus. The “*b*” arrows depict differences within the complex stimuli between local maxima and minima of the COVAS pain curves. Differences between complex and control stimuli (*a* arrows) and within-stimulus changes during complex stimuli (*b* arrows) were extracted for group-level analysis.

The parameters for the 3-step and constant, simple-step stimuli are similar to previously published reports investigating offset analgesia[7–9, 25–27]. The Inv and 2-step stimuli are novel stimuli with timing parameters and temperatures mirroring those of the 3-step stimulus. Temperature order of the Inv stimulus was manipulated to test whether an increase in temperature after a decrement produced a disproportionate increase in reported pain intensity. The 2-step stimulus is a modification of the 3-step stimulus to test whether eliminating the initial period at T1 affected offset analgesia magnitude. The 3-step was repeated in triplicate. Simple steps and Inv stimuli were repeated in duplicate. One 2-step was used. The number of replicates was chosen during preliminary testing. As noted above, the heat stimulus was sequentially rotated between three sites of the forearm. Three stimuli at each site (i.e. three rounds of three stimuli = 9 stimuli) was found to be feasible and acceptable to preliminary subjects (data not included). A T1 stimulus replicate was removed from the 9 stimuli and replaced with a 2-step stimulus. During data analysis, data from the T1 stimulus during the heat calibration procedure was included as a T1 replicate, allowing for the T1 condition to be done in duplicate. The order of these stimuli was randomized without replacement to ensure replicate testing using the randomization function in Excel (Microsoft, Redmond, Washington).

### Statistical Analysis

Replicate pain intensity curves were averaged within each subject. Since T1 was individually calibrated, pain intensity, thermode temperature, and time for each stimulus was sampled only during the timepoints at which thermode temperature was greater than T1-0.2°C. This epoch was further divided into t1 (5 seconds), t2 (5 seconds), and t3 (20 seconds) periods.

To identify extrema of pain intensity curves, the timepoint of transition from the t2 to t3 period was identified for each curve within each subject. Initially, the maximum or minimum during the 10-second epoch centered on the t2-to-t3 transition was calculated. Then, the subsequent minimum or maximum was determined in the time period following the initially calculated extremum. For example, in the analysis of the Inv pain intensity curve, the minimum during the 10-second epoch centered on the t2-to-t3 transition was obtained. The maximum following this minimum was subsequently obtained and recorded as the “local maximum” (e.g. Fig 4A). Similarly, in the analysis of the 2step pain intensity curve, the maximum during the 10-second epoch centered on the t2-to-t3 transition was first obtained, followed by measurement of the minimum value following this maximum (e.g. Fig 4E “local minimum”). This method was used to extract local pain intensity extrema during all stimuli, including during the subtracted pain intensity curves (Fig 5 and 6).

**Fig 2:**
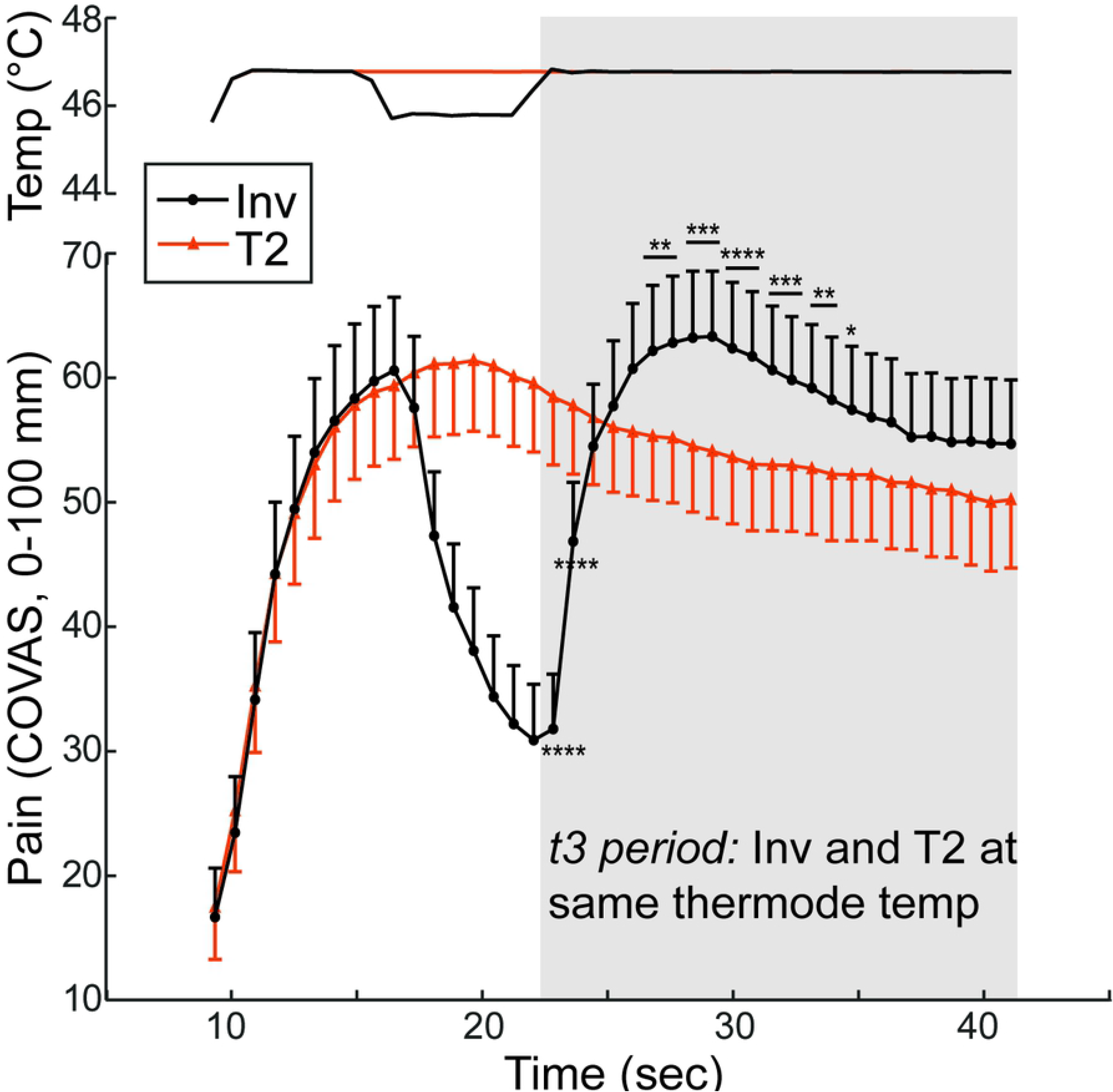
Pain intensity disproportionately increases following a transient, small decrease in temperature. **A.** Group mean temperature (top) and continuous pain intensity rating (bottom) curves from the Inv (black circles) and T2 control (orange triangles) stimuli are shown. Symbols represent group-level mean and error bars represent 95% confidence intervals. P-values: * p<0.05, ** p<0.01, *** p<0.001, **** p<0.0001.

**Fig 3:**
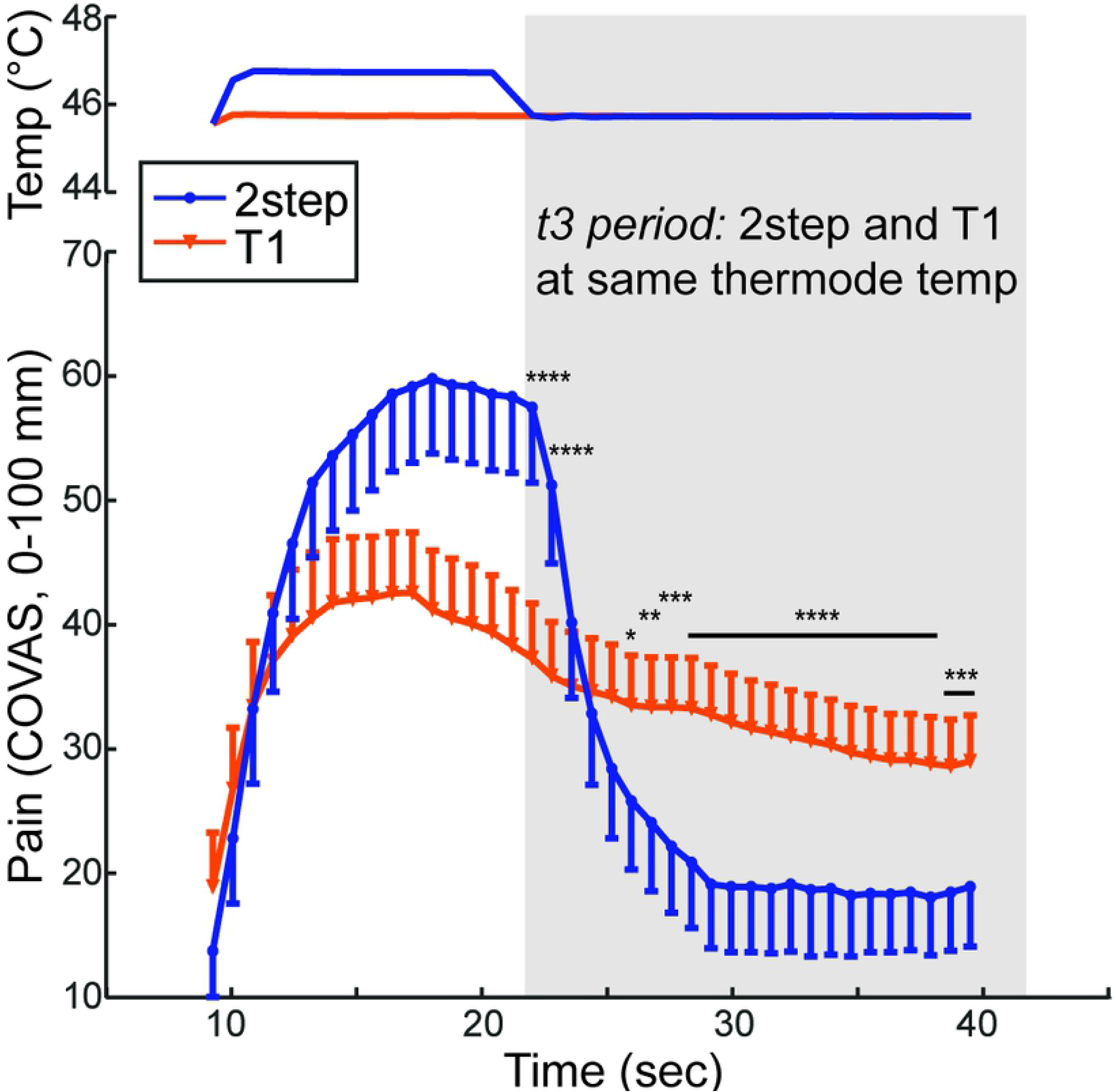
An isolated temperature decrease, without a preceding increase, reduces subsequent pain intensity. Group mean temperature (top) and continuous pain intensity rating (bottom) curves from the 2step (blue circles) and T1 control (orange triangles) stimuli are shown. Symbols represent group-level mean and error bars represent 95% confidence intervals. P-values: * p<0.05, ** p<0.01, *** p<0.001, **** p<0.0001.

**Fig 4:**
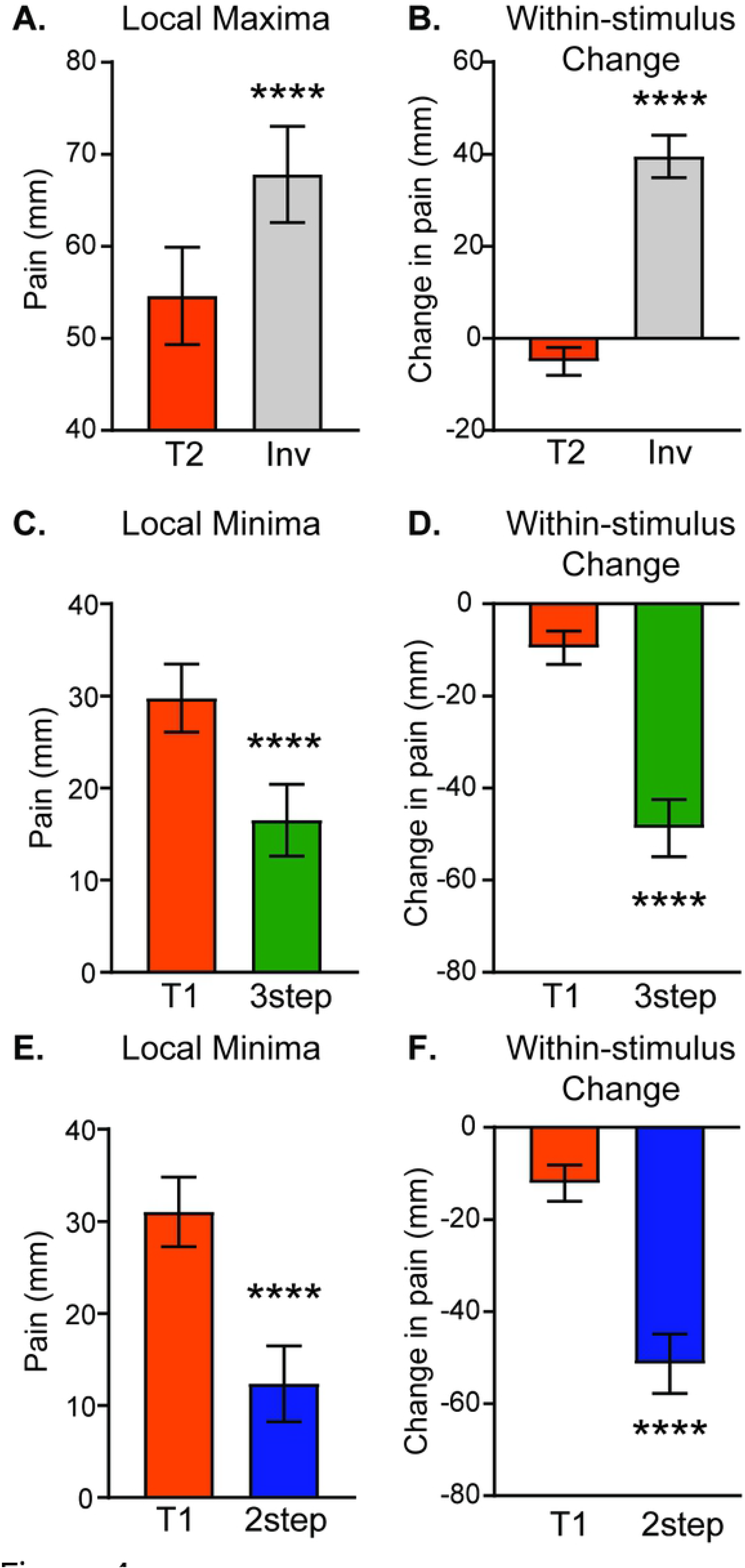
Extrema in continuously-rated pain intensity during complex stimuli are significantly different than controls. Pain intensity values on the computerized visual analogue scale were extracted at local maxima (A.&B.) and minima (C.-F.) for each subject as described in Fig 1. Pain intensities at matching timepoints were also extracted during constant control stimuli (T1 or T2). In the left column, pain intensities are compared between complex and control stimuli (arrow “a” in Fig 1). In the right column, the change in pain intensity within the complex curve is compared with the change in the control curve during the same time interval (arrow “b” in Fig 1). Group means with 95% CI are depicted in the bar graphs. Paired t-tests showed significant differences: **** p<0.0001.

**Fig 5:**
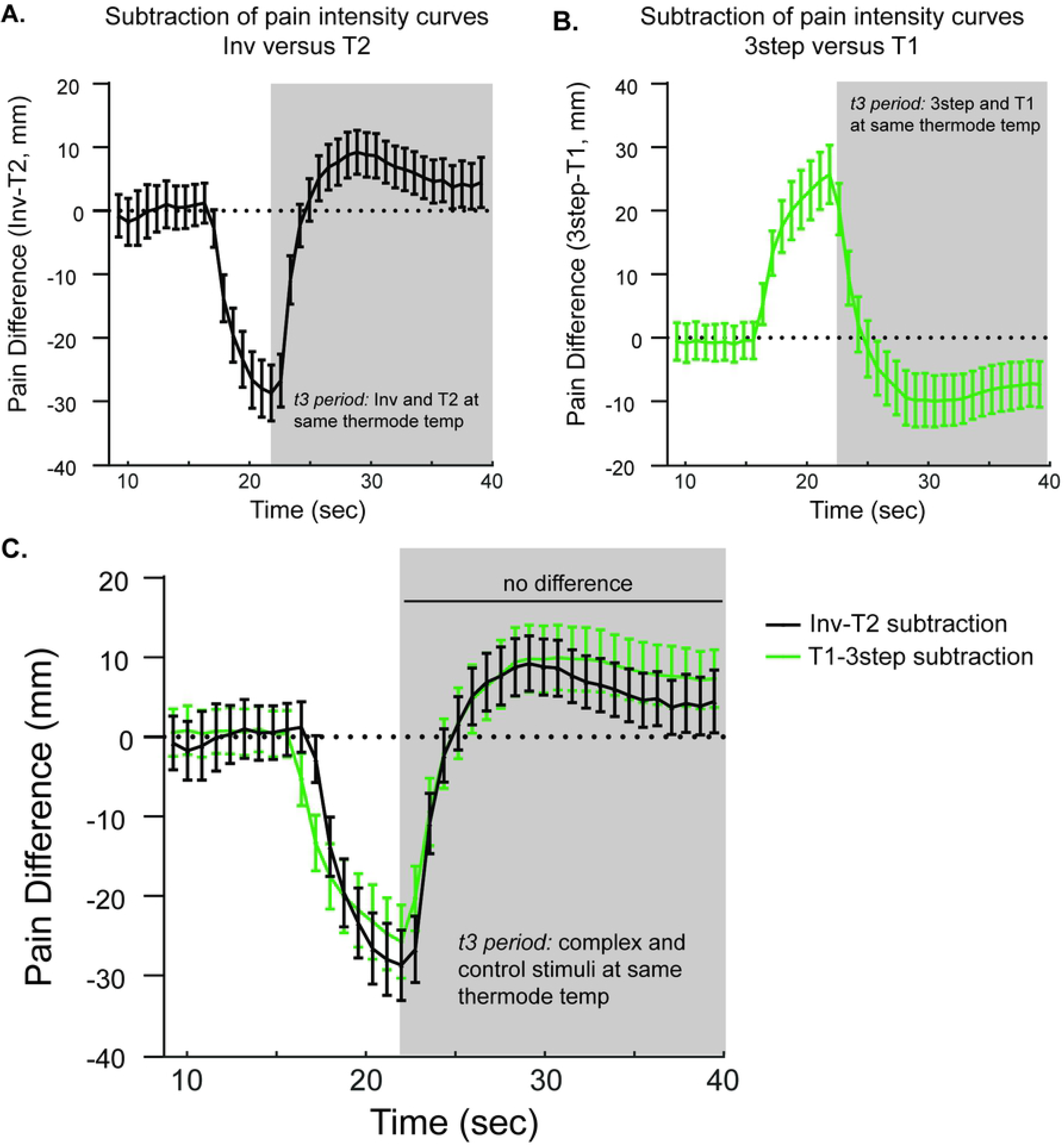
The magnitude of relative hyperalgesia and hypoalgesia with equivalent temperature increases and decreases is similar. **A.** Pain intensity curves during Inv and T2 stimuli were subtracted for each subject with group mean values plotted; error bars = 95% CI. **B.** Similarly, pain intensity curves during 3step and T1 stimuli were subtracted with group mean values plotted; error bars = 95% CI. In A. and B., shaded regions reflect the time period in which thermode temperatures are the same between complex (Inv or 3step) and constant (T2 or T1) stimuli. **C.** To compare pain difference curves, the inverse of the 3step-T1 subtraction curve in B. was plotted with the Inv-T2 subtraction curve. Means and 95% CI are shown at each timepoint.

**Fig 6:**
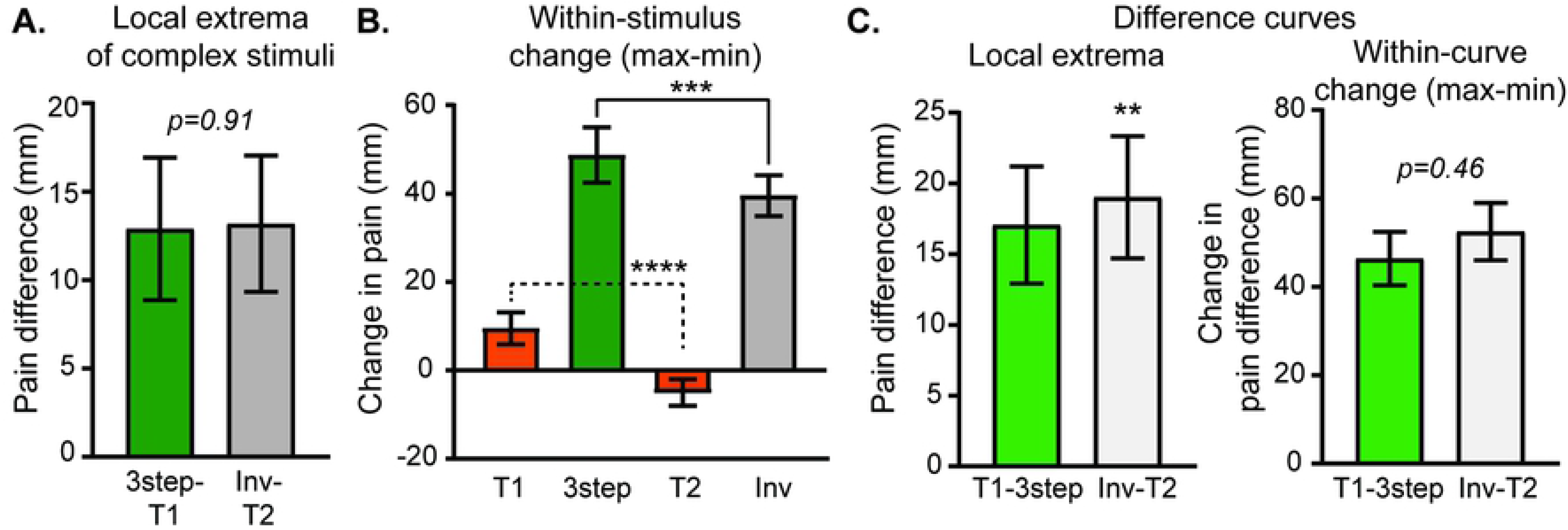
Within-subject analysis controlling for adaptation shows no significant difference in the magnitude of pain deviations in complex stimuli despite their opposite direction of change. **A.** The difference between pain intensity extrema (max for Inv, min for 3step) and the pain intensity during control stimuli (T2 for Inv, T1 for 3step), reflecting “a” arrows in Fig 1, was determined within each subject. **B.** Change in pain within stimulus (max – min, “b” arrows in Fig 1), for complex stimuli and matched timepoints during control stimuli. 1-way RM ANOVA showed significant main effect of stimulus. Post-hoc testing showed significant differences including those shown. **C.** Subtraction curves were analyzed within each subject for extrema and within-curve change. For all graphs, group mean with 95% CI error bars are plotted. P-values: ** p<0.01, *** p<0.001, **** p<0.0001.

For group-level analysis of pain intensity curves over time, pain intensity, thermode temperature and time were aligned so that the initial timepoint for all subjects occurred at the first timepoint where thermode temperature was greater than T1-0.2°C. The data from each subject was then downsampled to 1 Hz. Group mean and 95% confidence intervals were then calculated for each timepoint. To compare the pain intensity curves across stimuli, repeated measures 2-way ANOVAs were performed during the relevant time period with matching by stimulus and time with post-hoc testing using Sidak’s multiple comparisons test.

For group-level analysis of extrema, extrema were obtained from complex curves within each subject as outlined above and then the group-level mean and 95% confidence intervals were calculated. To compare with control curves, the timepoint of the extreme value was used to identify pain intensity at the same timepoint within each subject in the control curve (Fig 1 B-D, arrow “a”).

Within-stimulus change was analyzed by obtaining extrema and subtracting them within each subject. For example, for Inv, the local minimum was subtracted from the local maximum (Fig 1B, “b” arrow). The value of the subtraction was then averaged across the group to obtain the group-level mean. The change in pain intensity during the same timepoints as the complex pain intensity extrema was also extracted from the appropriate control pain intensity curves. For example, when comparing with the Inv stimulus pain intensity, pain intensity values from the T2 stimulus were obtained at the Inv pain intensity maximum and minimum. In this particular case, the direction of change is negative since the timepoint of the Inv maximum has a smaller pain intensity in the T2 stimulus than the timepoint of the Inv minimum, which is reflected in Fig 4B.

A similar approach was applied to extract extrema and determine within-curve changes for the difference curves. Again, these values were calculated within each subject, and then averaged for a group mean value.

For comparisons between stimuli extrema and within-stimulus/within-curve change, paired t-tests were performed. Given randomization of noxious stimulus order, these values were treated as independent observations, and therefore did not require additional nested analyses. Univariate correlations were assessed by calculating Pearson’s r coefficient and accompanying p-values. 95% CI of Pearson’s r were calculated with Fisher’s transformation. Univariate linear regressions were calculated to draw regression lines in Fig 9.

Subject data were collected and managed using REDCap hosted at UCSF [28]. Anonymized data from REDCap were exported as a flat file and combined with anonymized sensory testing data and survey data using Excel (Microsoft, Redmond, Washington). Graphical and statistical analysis was performed using Matlab R2016b (The MathWorks, Natick, Massachusetts), StataMP v14 (Statacorp, College Station, Texas) and Prism 7 (GraphPad Software, La Jolla, California).

## Results

### The onset of supra-threshold noxious heat disproportionately increases subjective pain intensity when preceded by a transient decrease in noxious heat

Healthy subjects (N=74; 35 female and 39 male, mean age of 28.2 years ± SD 7.2 years with range 18-50 years) underwent heat pain testing using a 30 mm by 30 mm thermode applied to the volar surface of the non-dominant forearm (Fig 1). The group mean heat pain threshold was 44.6 °C ± SD 1.7 °C. Ascending noxious-range 30-second heat steps were then used to calibrate the specific temperatures used in suprathreshold heat stimuli for each subject. The temperature that elicited a pain intensity rating of 50 mm at the end of a 30-second stimulus was defined as the T1 for an individual subject. The mean T1 used was 45.8°C ± SD 1.7°C ranging from 39°C to 47°C. T2 was defined as 1 C° hotter than T1. Using individualized T1 and T2 values, subjects underwent a battery of randomly-ordered suprathreshold heat stimuli, including both complex (Inv, 3step, and 2step) and control (T1 and T2) stimuli as outlined in Fig 1.

We tested the hypothesis that a rising cutaneous noxious heat stimulus produces a disproportionately greater reported pain intensity than when the same intensity stimulus is constant. Pain intensity ratings obtained during a novel complex suprathreshold heat stimulus, Inv, and a constant control stimulus, T2, were compared (Fig 2). During the t3 period in which thermode temperature is the same in both the Inv and control T2 stimuli (shaded area in Fig 2), reported pain intensity appeared greater in the Inv stimulus than the reported pain intensity in the control T2 stimulus. To understand the effect of stimulus type (Inv vs. T2) on reported pain intensity over time, a 2-way repeated-measures ANOVA during the t3 period with matching by stimulus and time was calculated and demonstrated a significant main effect of stimulus type (F (1.000, 73.00) = 6.041, p=0.0164), time (F (2.389, 174.4) = 23.27, p<0.0001), and time x stimulus type interaction (F (3.870, 282.5) = 46.54, p<0.0001). Sidak’s multiple comparison test confirmed statistically significant differences between Inv and control T2 stimuli at each timepoint noted in Fig 2 with asterisks reflecting corrected p-value thresholds. During the t3 period, despite identical thermode temperatures, reported pain intensity following a noxious stimulus increase in the Inv stimulus was significantly higher than the reported pain intensity during the constant T2 stimulus.

### The offset of supra-threshold noxious heat disproportionately decreases pain intensity report and does not require a preceding transient increase in noxious heat

To confirm the finding that small reductions in cutaneous noxious heat lead to disproportionate decreases in pain perception [8], pain intensity ratings during the complex suprathreshold heat stimulus frequently used to elicit offset analgesia (3step) were compared with pain intensity ratings during the control, constant T1 stimulus (S1 Fig). Using pain intensity data from the t3 period of 3step and T1 stimuli, a 2-way repeated-measures ANOVA with matching by stimulus and time demonstrated a significant main effect of stimulus type (F (1.000, 73.00) = 5.389, p=0.0231), time (F (1.899, 138.6) = 90.93, p<0.0001), and time x stimulus type interaction (F (2.718, 198.4) = 88.62, p<0.0001). Sidak’s multiple comparison test confirmed statistically significant differences noted in Supplemental Fig 1 with asterisks representing corrected p-value thresholds for between stimulus comparisons at each timepoint.

**Fig S1: S1 Fig: A transient increase then decrease in noxious heat decreases subsequent pain intensity.** Group mean temperature (top) and continuous pain intensity rating (bottom) curves from the 3step (green circles) and T1 control (orange triangles) stimuli are shown. Symbols represent group-level mean and error bars represent 95% confidence intervals. P-values: * p<0.05, ** p<0.01, *** p<0.001, **** p<0.0001.

To test whether the preceding transient increase to T2 is required for decreased pain intensity in the t3 period following a temperature offset, a two-step design was used (2step stimulus). Pain intensities during the 2step heat stimulus were compared with those during the constant control T1 stimulus (Fig 3). During the t3 period in which thermode temperature is the same, pain intensity during the 2step stimulus appears lower than the pain intensity during the T1 stimulus. This is a statistically significant difference, since a 2-way repeated-measures ANOVA with matching by stimulus and time showed a significant main effect of stimulus type (F (1.000, 73.00) = 15.16, p=0.0002), time (F (2.122, 154.9) = 62.01, p<0.0001), and time x stimulus type interaction (F (3.171, 231.5) = 55.36, p<0.0001) with significant *post hoc* testing using the Sidak multiple comparison test (corrected p-values denoted in Fig 3). Comparing pain intensity reported during the t3 period of either 3step or 2step with pain intensity during the same time period of the constant control stimulus T1 demonstrates that noxious range temperature decreases significantly decrease pain intensity.

### Within-subject analysis of pain intensity extrema confirms disproportionate changes in reported pain intensity with both noxious heat increases and decreases

To further characterize pain intensity differences between complex and control stimuli, local extrema were extracted from pain intensity curves during complex stimuli (Inv, 3step, and 2step) as discussed in the Methods section and Fig 1. For each planned comparison, the pain intensity at the same timepoint was extracted from the corresponding constant control curve (T1 or T2). Extrema and constant control pain intensity values were then averaged across the group to allow for group-level comparisons (Fig 1, arrow “a”). Additionally, within-stimulus pain intensity changes were calculated by subtracting the preceding minimum (for Inv) or maximum (for 3step and 2step) from the local extreme value in the t3 period (maximum for Inv and minimum for 3step and 2step; see Fig 1, arrow “b”). Both analyses have previously been used in studies of offset analgesia (e.g. [7, 9, 12, 25, 29, 30]).

Both analytic methods confirm that complex stimuli, with preceding changes in supra-threshold noxious heat, elicit significantly different pain intensities than constant control stimuli. The local maximum of the reported pain intensity curve during the Inv stimulus is significantly greater than pain intensity during the T2 stimulus at the equivalent timepoint (Fig 4A). Additionally, the within-stimulus pain intensity change (Fig 1B, arrow “b”) is significantly greater during the Inv stimulus than the change measured at equivalent timepoints during the constant T2 stimulus (Fig 4B). Notably, the direction of change is different between the Inv and T2 stimuli. The T2 stimulus has a negative change, reflecting a decrease in pain intensity. The pain intensity difference at the local maximum of the Inv stimulus (Fig 4A) is 13.2 mm (95% CI 9.3 mm – 17 mm), which appears to be greater in absolute magnitude than an estimate of offset analgesia magnitude calculated in a recent quantitative meta-analysis of −4.6 mm (95% CI −7.5 mm – – 1.7 mm)[27].

For noxious heat decreases, the pain intensity minima during both 3step and 2step stimuli (Fig 4C and 4E respectively) were significantly lower than equivalent timepoints (uniquely determined for 3step and 2step) during the control T1 stimulus. Similarly, the change in pain intensity within each stimulus was more negative, reflecting decreasing pain intensity, than that observed in the control T1 stimulus (Fig 4D and 4F).

### Comparison of Inv and 3step stimuli shows a similar absolute magnitude of changes in pain intensity despite opposite direction of change

To account for time-dependent changes in pain intensity, such as adaptation, and allow for comparison across complex stimuli, pain intensity curves during constant control stimuli, T2 and T1, were subtracted from pain intensity curves during complex stimuli, Inv and 3step respectively. This was done within each subject and then averaged at each timepoint across the group. The Inv-T2 pain difference curve with 95% confidence intervals is plotted in Fig 5A, and the 3step-T1 pain difference curve is plotted in Fig 5B. To compare their relative absolute magnitude, the 3step-T1 pain difference was inverted and plotted on the same axis as the Inv-T2 pain difference curve (Fig 5C). Graphically, there appears to be substantial overlap between these two curves. During the t3 period, a repeated measures 2-way ANOVA with matching by both factors (subtraction curve and time) only showed a main effect of time (F (2.985, 217.9) = 83.84, p<0.0001) but no effect of subtraction curve type (F (1.000, 73.00) = 0.8225, p=0.367) or time by curve type interaction (F (3.785, 276.3) = 2.440, p=0.051). Post-hoc testing showed no significant differences between the Inv-T2 and T1-3step subtraction curves at individual timepoints (Sidak’s MCT). Overall, these results suggest that increases and decreases in temperature at the t2-to-t3 transition produce changes in pain intensity of opposite sign but highly similar absolute magnitude.

Pain intensity reported during the complex stimuli was further compared between Inv and 3step stimuli using extrema and within-stimulus change analysis described above and in Fig 1. In Fig 6A, the absolute magnitude of the difference between complex stimulus extrema (minimum for 3step and maximum for Inv) and the equivalent timepoint in the control stimulus (T1 for 3step and T2 for Inv; Fig 1, “a” arrows) was averaged across the group and plotted with 95% confidence intervals. A paired t-test shows no significant difference between these two values. In Fig 6B, the group mean average of within-stimulus change (Fig 1, “b” arrows), maximum – minimum for 3step with equivalent timepoints during T1 and maximum – minimum for Inv and equivalent timepoints during T2, are plotted. To compare within-stimulus pain intensity change across stimulus type, a repeated measures 1-way ANOVA with matching by subject was calculated and showed a significant main effect of stimulus type (F (1.901, 138.8) = 149.7, p<0.0001). Planned post-hoc testing with Sidak’s MCT showed significant differences not only between 3step and Inv, but also between the constant control stimuli (T1 versus T2). Interestingly, the mean differences between Inv and T2 (44.5 mm; 95% CI 36.9 mm-52.1 mm) and 3step and T1 (39.2 mm; 95% CI 32.0 mm-46.4 mm) appeared similar, as did the mean differences between Inv and 3step (9.2 mm; 95% CI 3.7 mm-14.6 mm) and T2 and T1 (14.5 mm; 95% CI 8.1 mm-21.0 mm). This suggests that the observed difference between Inv and 3step in Fig 6B may actually be due to time-dependent changes shared across all stimuli (e.g. adaptation). To account for this possibility, the difference curves (Inv-T2 and T1-3step, plotted in Fig 5C) were analyzed. Within each subject, the maxima of the difference curves following the minima occurring around the t2-to-t3 transition was determined and then averaged across the group. Fig 6C (left) shows that the maxima of the Inv-T2 curve during the t3 period was in fact slightly larger than the maxima of the T1-3step (paired t-test; mean difference 6.1 mm; 95% CI 2.1 mm – 10.1 mm). However, the within-curve change of the subtraction curves did not reach a statistically significant difference. Taken together, it appears that the increase in pain intensity following the noxious heat increase during the Inv stimulus is similar in magnitude to the decrease in pain intensity following the noxious heat decrease during the 3step stimulus.

### A simple noxious stimulus increase, if not preceded by a decrement, produces a much smaller increase in pain intensity

The effect of the temperature increase at the t1-to-t2 transition on subsequent pain intensity during the t2 period was also explored. During the t2 period, the thermode temperatures are the same between 3step, T2 and the 2step stimuli. Therefore, comparisons between 3step and either T2 or 2step stimuli should inform the effect of a simple temperature increase. Group averaged time series data of pain intensity are plotted in Fig 7A and 7C. Group averaged local maxima during the 3step stimulus are compared with pain intensity values during control stimuli, T2 (Fig 7B) and 2step (Fig 7D). There were small differences between the 3step and controls that reached statistical significance with paired t-tests. Repeated measures 2-way ANOVAs during the t2 period were calculated. For 3step versus T2 (Fig 7A), there were main effects of time (F (1.634, 119.3) = 103.9, p<0.0001), stimulus (F (1.000, 73.00) = 6.801, p=0.0110), and time x stimulus (F (1.790, 130.7) = 99.69, p<0.0001). For 3step versus 2step, there were main effects of time (F (1.422, 103.8) = 58.23, p<0.0001) and time x stimulus (F (1.594, 116.4) = 63.69, p<0.0001), but no main effect of stimulus (F (1.000, 73.00) = 1.364, p=0.2466). Post-hoc testing using Sidak’s MCT showed significant differences labeled on the graphs with asterisks. Although apparent only with analysis of pain intensity curve maxima, there appeared to be a small increase in pain intensity during the t2 period of the 3step stimulus compared with control stimuli.

**Fig 7:**
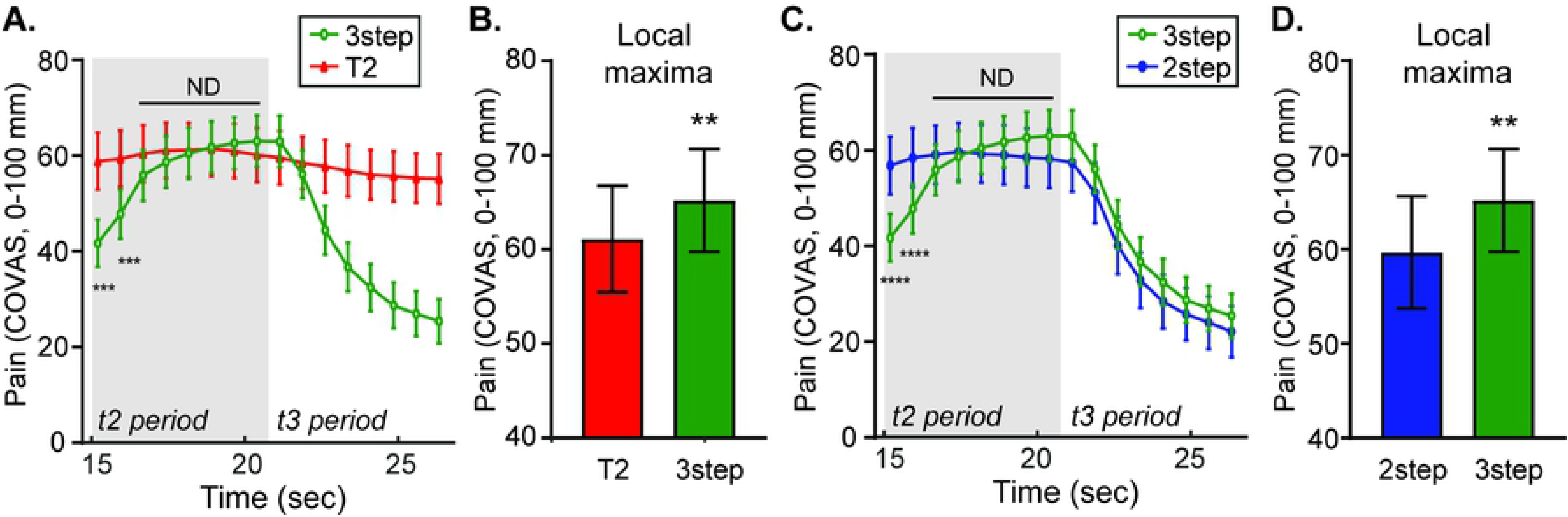
An increase in temperature within the noxious range without an immediately preceding decrease produces a small increase in pain intensity. **A.** Group mean continuous pain intensity rating curves from the 3step (green circles) and T2 control (red triangles) stimuli are plotted. **B.** Group mean pain intensity local maxima during the 3step stimulus and the pain intensity at the equivalent timepoint during the control T2 stimulus are plotted. **C.** Group mean continuous pain intensity rating curves from the 3step (green circles) and 2step (blue circles) stimuli are plotted. **D.** Group mean pain intensity local maxima during the 3step stimulus and the pain intensity at the equivalent timepoint during the control 2step stimulus are plotted. For all graphs (A.-D.), group means are plotted with error bars representing 95% CI. P-values: ** p<0.01, **** p<0.0001. ND = no difference (p>0.05).

### A simple decrease in noxious heat elicits a subtly larger offset analgesia than a decrease preceded by an increase

Comparing pain intensity during 3step and 2step stimuli interrogates the perceptual effect of a prior temperature increase on a subsequent decrease in the t3 period. Group mean timeseries data are plotted in Fig 8A with comparisons of local minima, within-stimulus change, and pain difference analyses shown in Fig 8B-D. Comparing pain intensity at the local minima and the difference between the local minima the constant control stimulus T1 (Fig 1C&D, “a” arrows) shows a slightly larger magnitude offset analgesia (more negative) in the 2step stimulus than the 3step stimulus that reaches statistical significance (Fig 8B, paired t-tests). Fig 8B also shows there is no difference when comparing within-stimulus changes in pain intensity between 2step and 3step stimuli (Fig 1C&D, “b” arrows). A repeated measures 2-way ANOVA comparing pain intensity curves during the t3 period of 2step and 3step stimuli (Fig 8) revealed a main effect of time (F (2.325, 169.7) = 102.8, p<0.0001) and stimulus type (F (1.000, 73.00) = 8.518, p=0.0047) without an interaction (F (3.852, 281.2) = 0.4906, p=0.7357). Post-hoc testing with Sidak’s MCT showed no single timepoint difference achieved statistical significance. Taken together, it appears that offset analgesia is slightly larger in the 2step stimulus compared with the 3step stimulus.

**Fig 8:**
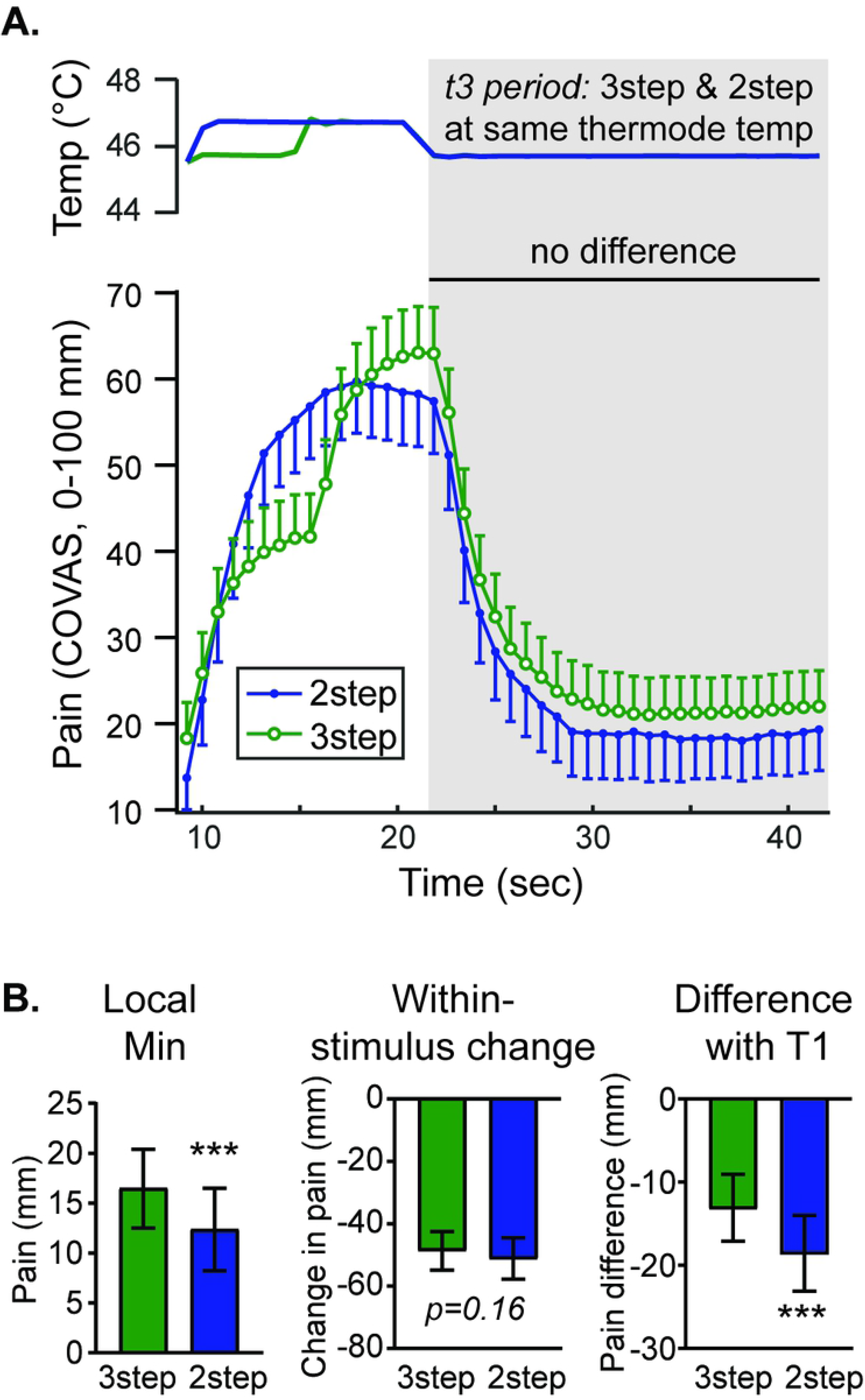
A noxious temperature decrease without a prior increase produces a subtly larger decrease in pain intensity. **A.** Group mean temperature (top) and continuous pain intensity rating (bottom) curves from the 2step (blue circles) and 3step (green circles) stimuli are shown. Symbols represent group-level mean and error bars represent 95% confidence intervals. Although a 2-way RM ANOVA with matching by stimulus and time showed main effects of time and stimulus, there was no statistically significant difference at any timepoint during the t3 interval. **B.** Group means obtained by within-subject analysis of minima (local min), the change in pain from maxima to minima (within-stimulus change) and the difference between complex curve minima and T1 at equivalent timepoints (difference with T1) are plotted with error bars representing 95% CI. P-values: *** p<0.001.

### Although stimulus temperature is not correlated with disproportionate decreases or increases in pain during 3step and Inv stimuli, the magnitudes of the two are inversely correlated

In the initial characterization of offset analgesia in 12 volunteers, the magnitude of offset analgesia was consistent across a range of noxious temperatures [8]. In the current larger dataset, there again appears to be no correlation between initial temperature and offset analgesia (Fig. 9A&B, Table 1). No correlation exists between offset analgesia (measured as the local minimum during the 3-step stimulus minus the pain rating during the constant T1 stimulus, Fig 1C “a” arrow) and either heat pain threshold or T1 temperature (Fig. 9A&B; R^2^<0.0001 and R^2^= 0.0005 respectively). This is consistent with other studies, in which offset analgesia was demonstrated following heat stimulus changes from a range of initial temperatures. Additionally, there is no correlation between pain intensity amplification during the Inv stimulus (measured as the local maximum during the Inv stimulus minus the pain rating at the same time point during the constant T2 stimulus, Fig 1B “a” arrow) and either heat pain threshold or T1 temperature (Fig. 9C&D, Table 1; R^2^=0.0006 and R^2^= 0.007 respectively). Using other measures of perceptual enhancement of temperature changes, including within-stimulus change (Fig 1B&C, “b” arrows), local extrema of subtraction curves (Fig 6C), and within-curve change of subtraction curves (Fig 6C), again there was generally no correlation with stimulus temperature or other variables listed above (Table 1). We did find a weak correlation between Inv within-stimulus change (Table 1; Fig 1B, “b” arrows) and both T1 temperature used (R^2^=0.0762) and heat pain threshold (R^2^=0.0538), but this was not seen with other measures of perceptual enhancement. Taken together, it appears that there is minimal relationship between the magnitude of the perceptual amplification or inhibition of pain produced by small temperature changes and the initial noxious stimulus intensity prior to the change.

**Fig 9:**
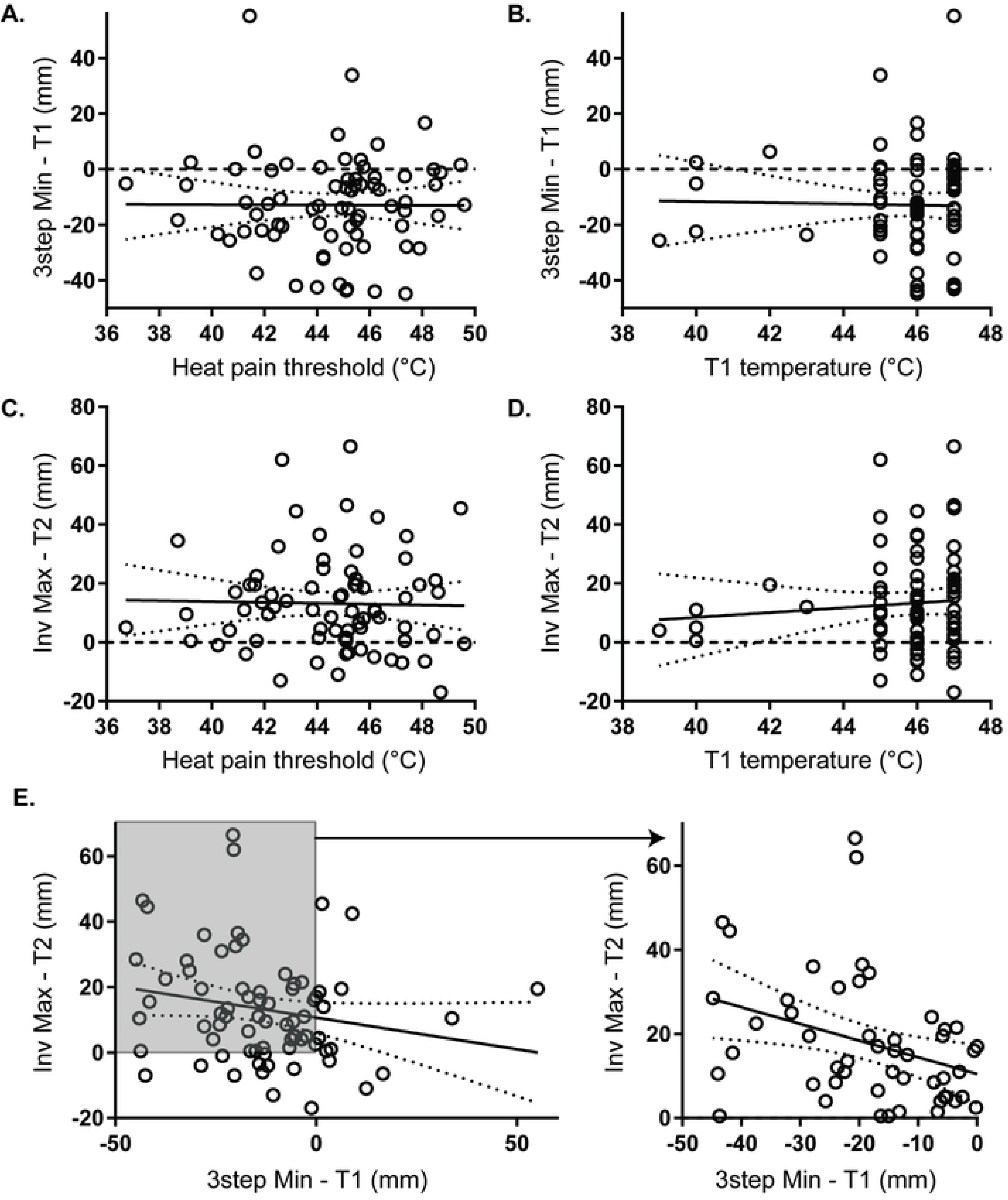
Pain intensity amplification with either increases or decreases are independent of temperature but are correlated with each other. Scatter plots are shown with each point representing an individual subject. Solid lines represent best-fit lines from linear regression analysis. Dotted lines represent bounds of the 95% confidence intervals calculated as part of the linear regression. No correlation exists between temperatures (heat pain threshold (A. or C.) or T1 stimulus temperature used (B. or D.)) and pain intensity amplification with noxious temperature decreases (3step Min-T1) and increases (Inv Max-T2). There is a trend toward an inverse correlation between perceptual enhancement of increases and decreases (E.) which becomes significant in the subgroup of subjects in grey (N=50, R^2^=0.12, p=0.014).

**Table 1: Correlation coefficients between measures of perceptual enhancement of noxious stimulus increases, offset analgesia, and psychosocial attributes.**

As an initial comparison between perceptual enhancement of noxious stimulus increases and decreases, pairwise correlations were made using measures outlined above. Interestingly, there were moderate to strong inverse correlations observed using measures of within-stimulus change (Table 1; r = 0.73, 95% CI 0.61 – 0.82; Fig 1B&C, “b” arrows) and the within-subtraction-curve change (Table 1; r = 0.80, 95% CI 0.70 – 0.87; Fig 6C, right). A weak inverse correlation was observed using subtraction curve extrema (Table 1; r = 0.25, 95% CI 0.017 – 0.448). Using the difference between complex stimuli extrema and control stimuli (Fig 1B&C, “a” arrows), there was no significant correlation between pain intensity amplification following increases and decreases, although there appeared to be a trend towards significance (Table 1 and Fig 9E; r = −0.22, 95% CI −0.416 – 0.022; R^2^=0.041, deviation of slope from zero: p=0.084). In a subset of subjects who had both offset analgesia (negative values) and increased pain intensity with temperature increases in Inv versus T2 (positive values), a linear regression did reveal a significant correlation (Fig 9E; linear regression, N=50, R^2^= 0.12, deviation of slope from zero: p=0.014).

Additionally, neither offset analgesia nor the disproportionate increase in pain intensity during the Inv stimulus correlated well with age, sex, body mass index, socioeconomic status, self-report measures of pain catastrophizing, depression, anxiety, or impulsivity (univariate correlations; Table 1). There were a few weak correlations that did achieve statistical significance that are noted in Table 1. Overall, there does appear to be a significant correlation between offset analgesia and perceptual enhancement of noxious stimulus increases.

### Discussion

The disproportionate drop in subjective pain sensation elicited by a decreasing noxious stimulus has been termed offset analgesia (OA) [7, 8]. Here, we demonstrate a disproportionate enhancement when the noxious stimulus is increasing (onset hyperalgesia, OH). Several lines of evidence support the existence of OH found in our study. First, using a novel heat stimulus in which a transient decrease in temperature from T2 to T1 is followed by a return to T2 (Inv stimulus), we observed significantly elevated pain intensity ratings at T2 compared with those reported in the constant control T2 stimulus. This was apparent using group-mean time series data (Fig 2) as well as within-subject analysis of local maxima and change in pain (Fig 4A&B). Additionally, comparing complex curves (Inv and 3step) showed similar perceptual enhancement with temperature increases (Inv) and decreases (3step). The InvT2 subtraction was no different from the sign-inverted 3stepT1 subtraction (Fig 5). Two different analyses also showed high similarity between OA and OH time course and magnitude measured as both difference from complex curve extrema and within-stimulus changes (Fig 6). Consistent with the known influence of prior pain experience on current pain intensity, we observe that the magnitude of both OA and OH is affected by the prior trajectory of the noxious stimulus. Finally, the magnitude of OA has a moderate inverse correlation with that of OH. Overall, these results are consistent with a unifying model of OA and OH in which the direction of change of noxious stimulus intensity strongly influences pain perception.

Prior studies support the notion of OH. Using radiant heat with an infrared laser, Morch and colleagues demonstrated that there is perceptual enhancement of temperature increases compared with a constant stimulus[15]. However, their model predicting pain intensity showed larger magnitude perceptual enhancement with temperature decreases than increases. The current study extends these findings by reporting the first evidence of OH using contact heat and demonstrates similar absolute magnitudes of OA and OH. Additionally, our findings provide empirical support for a non-linear model of pain intensity that incorporates perceptual feedback proposed by Apkarian and colleagues [14]. This model predicted a rapid increase in pain intensity observed with the increase from T1 to T2 during the t2 period of a 3step stimulus. The authors noted the change to be more subtle than the OA effect, but did not further quantify it. We quantified this small increase in pain intensity by comparing the 3step stimulus (Fig 7) with the control T2 stimulus. Importantly, our observation of different magnitudes of OH and OA depending on the prior trajectory of the heat stimulus is consistent with a perceptual feedback model incorporating change in noxious stimulus intensity as a variable that predicts subjective pain intensity. Overall, the observation that the novel noxious heat stimulus, Inv, elicits onset hyperalgesia complements and extends prior work suggestive of the phenomenon.

We also found evidence of OA following a simple noxious heat decrease in the 2step stimulus that was equivalent if not subtly larger than OA measured during the 3step stimulus. In contrast, Haggard and colleagues recently reported no changes in pain intensity with isolated temperature decreases compared with constant stimuli[31]. This difference may be due to technical reasons including the use of individually tailored stimulus temperatures in the current study as opposed to predetermined temperatures and the possible confound of concurrent mechanical stimulation with Haggard and colleagues’ thermode setup. Haggard and colleagues suggest an alternative reason, in which a certain duration in the noxious range is required to elicit OA. Kurata and colleagues reported greater magnitude OA with a longer T2 duration following a constant, 5-second T1 duration in a three-step stimulus design[32]. Taken together with our observation of OA in the 2step stimulus, we suggest Haggard and colleagues would have found OA and OH with longer-duration noxious stimulation prior to the simple temperature changes.

Although identifying neural mechanisms of OH is beyond the scope of the current study, we did find similarity with OA, which is thought to involve central processing based on several behavioral and neuroimaging experiments[9, 10, 14, 26, 33, 34]. Like OA[8], the temperature of T1 used does not correlate well with the magnitude of OH (Fig 9 and Table 1). Heat pain threshold, which was measured prior to supra-threshold testing, also does not correlate well with either OA or OH. Interestingly, there may be a correlation between OA and OH, although the strength of correlation depends on how each is measured. Within-stimulus change in pain intensity (Fig 1 “b” arrows) and within-curve change of the subtraction curves show moderate correlations between OH and OA. Analysis of extrema (Fig 1B “a” arrows) show a weak correlation between OH and OA, which becomes moderate in the subgroup of subjects with both OA and OH (Fig 9E). Interestingly, within-stimulus change measures show only weak-moderate correlation with extrema difference measures (Fig 1B “b” versus “a” arrows; Table 1), consistent with divergent measurement effects observed in a recent meta-analysis of OA[27]. Given the stronger correlation between OA and OH using the within-stimulus change, it seems possible that studies only analyzing that measure may be capturing a composite outcome reflecting both OA and OH[8, 9, 13, 25, 33, 35–37]. Certainly, additional studies are required to clarify the mechanisms of OH and its relationship to OA in different populations.

Given our findings of OH and subtle differences in magnitude of both OH and OA depending on immediately prior noxious stimulus intensity, we favor an explanatory model similar to that proposed by Apkarian and colleagues[14] and outlined in [38] whereby changing noxious stimulus intensity impacts predictions of pain and pain relief which modulates nociceptive transmission and pain perception bidirectionally. We observed that the magnitude of OH was larger during the Inv stimulus than during the 3step stimulus. This could be due to differences in how predicted pain intensity changes, which will differ between these two stimuli. According to our proposed model, the transient drop during the t2 period in the Inv stimulus indicates that future stimulus intensity will likely decrease, reducing the motivation to engage in an action to terminate the noxious input. The rise back after the earlier decrease reverses the direction of the prediction and increasing the motivation to respond to the noxious stimulus. According to the Motivation-Decision model [39], this switch in action selection will engage a top down modulatory circuit that amplifies pain if the decision is to respond to it and inhibits pain if the decision is to ignore the pain. In different behavioral paradigms, changed predictions/expectations about pain intensity can elicit either increases or decreases in reported pain [6, 40, 41].

On the other hand, the magnitude of OA was only subtly greater during the 2step stimulus as compared to the 3step stimulus. This could still be due to within-stimulus predictions, since the 2step stimulus did not have an initial period at T1, but only included a temperature decrease from T2 to T1. It is possible that the difference in perceptual enhancement between the stimuli is smaller than that observed between the Inv and 3step stimuli because the trajectory of pain intensity was generally the same – a noxious stimulus increase followed by a decrease. This combined model of OA and OH magnitude is supported by prior theoretical work[14] and by the observation that longer durations of the t2 interval in the three-step paradigm produce larger magnitude offset analgesia[32], since predictions can be time-dependent and continued pain predicts subsequent pain.

Alternative explanations for our observation of OH remain possible. Although our study design controls for time-dependent within-stimulus changes in pain intensity, such as adaptation or habituation, by incorporating constant control stimuli (T1 and T2), it is possible that there are independent competing time-dependent processes producing the pain intensity curves. For example, the increase in pain intensity in the t3 period of the Inv stimulus may result from temporal summation which is distinct from pain inhibiting processes, such as adaptation or OA. We do not favor this explanation since heat pain temporal summation occurs with more frequent temperature changes of at least 0.33 Hz and not at 0.25 Hz or less[42], which is in the range of the Inv stimulus. Alternatively, the temperature decrement in the Inv stimulus during the t2 interval may reverse a single pain inhibiting process, reflecting a dishabituation. The current study cannot rule this possibility out, but given the known bidirectional effects of expectancy/predictions on pain intensity and previous reports supportive of OH without the temperature decrement[14, 15], we favor the motivation-decision model. Future studies designed to manipulate pain and pain relief predictions will help delineate mechanisms on the behavioral level.

In conclusion: The current study establishes the existence of OH and posits a model of OA and OH emphasizing how the predictive nature of changes in pain intensity strongly influencing individual responses to such changes through top down bidirectional modulation of nociceptive transmission.

## Acknowledgements

The authors acknowledge Nidhi Anamkath for administrative and technical support.

